# Proximity based proteomics reveals Git1 as a regulator of Smoothened signaling

**DOI:** 10.1101/2025.01.06.631593

**Authors:** Jingyi Zhang, Gurleen Kaur, Eva Cai, Oscar Torres Gutierrez, Xiaoliang Liu, Sabyasachi Baboo, Jolene K Diedrich, Ju-Fen Zhu, Benjamin R. Myers, John R Yates, Xuecai Ge

## Abstract

The GPCR-like protein Smoothened (Smo) plays a pivotal role in the Hedgehog (Hh) pathway. To initiate Hh signaling, active Smo binds to and inhibits the catalytic subunit of PKA in the primary cilium, a process facilitated by G protein-coupled receptor kinase 2 (Grk2). However, the precise regulatory mechanisms underlying this process, as well as the events preceding and following Smo activation, remain poorly understood. To address this question, we leveraged the proximity labeling tool TurboID and conducted a time-resolved proteomic study of Smo-associated proteins over the course of Hh signaling activation. Our results not only confirmed previously reported Smo interactors but also uncovered new Smo-associated proteins. We characterized one of these new Smo interactors, Grk-interacting protein 1 (Git1), previously known to modulate GPCR signaling. We found that Git1 localizes to the base of the primary cilium, where it controls the cilium transport of Grk2, an early event in Hh signaling. Loss of Git1 impairs Smo phosphorylation by Grk2, a critical step for Smo-PKA interaction, leading to attenuated Hh signaling and reduced cell proliferation in granule neuron precursors. These results revealed a critical regulatory mechanism of Grk2 phosphorylation on Smo in the primary cilium. Our Smo-TurboID proteomic dataset provides a unique resource for investigating Smo regulations across different stages of Hh pathway activation.

## INTRODUCTION

The Hedgehog (Hh) signaling pathway is one of the most fundamental and highly conserved mechanisms governing embryonic development and tissue homeostasis. Its precise regulation orchestrates the formation of nearly every organ in vertebrates^1–3^. Insufficient Hh signaling leads to birth defects, ranging from holoprosencephaly to cardiovascular abnormalities and skeletal malformations^4–6^. Conversely, aberrant activation of Hh signaling drives the development of aggressive cancers, including medulloblastoma, a malignant pediatric brain tumor, and basal cell carcinoma, a common form of skin cancer^7–9^. Understanding the transduction mechanism of Hh signaling is essential for elucidating the underlying causes of Hh-related disorders.

At the core of Hh pathway is Smoothened (Smo), an atypical G protein-coupled receptor that transduces the signal across the cell membrane, ultimately triggering transcription of Hh target genes in the nucleus. In the absence of Sonic Hedgehog (Shh), Smo activity is inhibited by Patched (Ptch) which restricts Smo access to sterols, the Smo endogenous agonists^10–12^. Protein Kinase A (PKA) and Suppressor of Fused (SuFu), negative regulators of Hh pathway, inhibit Glioma-associated (Gli) transcription factors, thereby blocking Hh signaling^13–15^. Upon Shh binding to Ptch, Ptch-mediated inhibition is lifted, allowing Smo to be activated and accumulate in the primary cilium. How Smo transmits its signal to the downstream transducers has been an enigma in the field. Recent discoveries from our and collaborators’ lab offer significant insights into this process. Smo, via a PKA pseudosubstrate motif in its cytoplasmic domain, directly binds to the catalytic subunit of PKA (PKA-C), thereby inhibiting PKA activity^16,17^. The Smo-PKA interaction is facilitated by G protein-coupled receptor kinase 2 (Grk2), which phosphorylates the cytoplasmic tail of active Smo^18^. Genetic and pharmacological studies revealed that Grk2 phosphorylation on Smo is required for Smo-PKA interaction in the cilium^19^.

While these studies established the Smo-PKA interaction as the key event in Smo signaling, and identified Grk2 phosphorylation of Smo as a critical step that facilitates the recruitment of PKA-C, several questions concerning this key signaling process remain unanswered. First, our previous work demonstrated that Grk2 can directly phosphorylate active Smo in the cilium^19^. However, direct evidence showing that ciliary Grk2 is the sole source of Smo phosphorylation is missing. As Grk2 also has been shown to localize to the basal body^20^, it remains unclear whether ciliary or basal body-localized Grk2 serves as the primary source of Smo phosphorylation. Second, we previously showed that Grk2 enters the primary cilium immediately following Hh pathway activation, but what modulates the ciliary translocation of Grk2 remains unclear. Finally, Smo accumulates in the cilium after Hh pathway activation, a process mediated by its interactions with a variety of molecules. The identity of proteins specifically involved in Smo trafficking and signaling at different stages of Hh signal transduction remains unclear.

To address these questions, we took the approach of systematically mapping Smo interactions across different stages of Hh signal transduction. Previous efforts to identify Smo interactors have employed co-immunoprecipitation (Co-IP) based strategies^21^, which are not ideal for capturing transient or low-abundance interactions. To overcome this limitation, we employed TurboID, a proximity-based biotin ligase with improved labeling kinetics over its precursor BioID^22^. In a previous study, we successfully used TurboID to identify new ciliary proteins that regulate Hh signaling^23^, demonstrating that the labeling kinetics of TuboID enable capture of the often-transient protein interactions essential for signal transduction. Here, we fused TurboID to the cytoplasmic tail of Smo and selected the Smo-TurboID stable cell line with near-physiological levels of Smo expression. From time-resolved proteomics with the Smo-TurboID stable cell line, we discovered distinct cohorts of Smo-associated proteins at different stages of Hh signaling. Among them are known Smo interactors, such as PKA-C, Dlg5, Evc2, Grk2, and Wdr35 ^24–26^, as well as a group of new Smo-associated proteins. We characterized one of these new candidates, Grk-interacting protein (Git1), a multidomain protein that acts as GTPase-activating protein for ADP ribosylation factor (ArfGAP)^27^. Git1 loss in mice leads to defects in pulmonary vascular formation and microcephaly-like phenotypes^28–30^, consistent with impaired Hh signaling transduction. We found that Git1 localizes to the base of the primary cilium and is required for Grk2 translocation into the cilium at the initial stage of Hh pathway activation. Git1 knockout has no impact on Grk2 levels at the basal body, but abolishes Grk2 translocation to the cilium. This subsequently reduces Smo phosphorylation in the cilium and diminishes Hh signaling. Finally, selectively targeting Grk2 to the cilium is sufficient to restore Hh signaling in Git1 knockout cells.

Our findings highlight Git1 as a critical regulator for Grk2 phosphorylation of Smo, and pinpoint the cilium as the primary site where this phosphorylation occurs. Broadly, our time-resolved proteomic dataset with Smo-TurboID provides a valuable resource for understanding the precise regulation of Smo signaling during Hh pathway activation.

## RESULTS

### Generating the Smo-TurboID stable cell line to mimic endogenous Smo during Hh signaling activation

To systematically identify transient and weak interactions of Smoothened (Smo), we leveraged the proximity labeling tool TurboID. We fused the V5 epitope and TurboID to the C-terminal cytoplasmic tail of mouse Smo (Fig 1A), which enables the biotinylation of proteins within a 10-20 nm radius of Smo. Upon the Shh ligand stimulation, Smo translocates from the extraciliary regions to the cilium^31,32^. This allows the capture of biotinylated proteins at various time points during Hedgehog (Hh) signaling activation, and provide snapshots of Smo-interacting proteins throughout this process.

**Figure 1.**
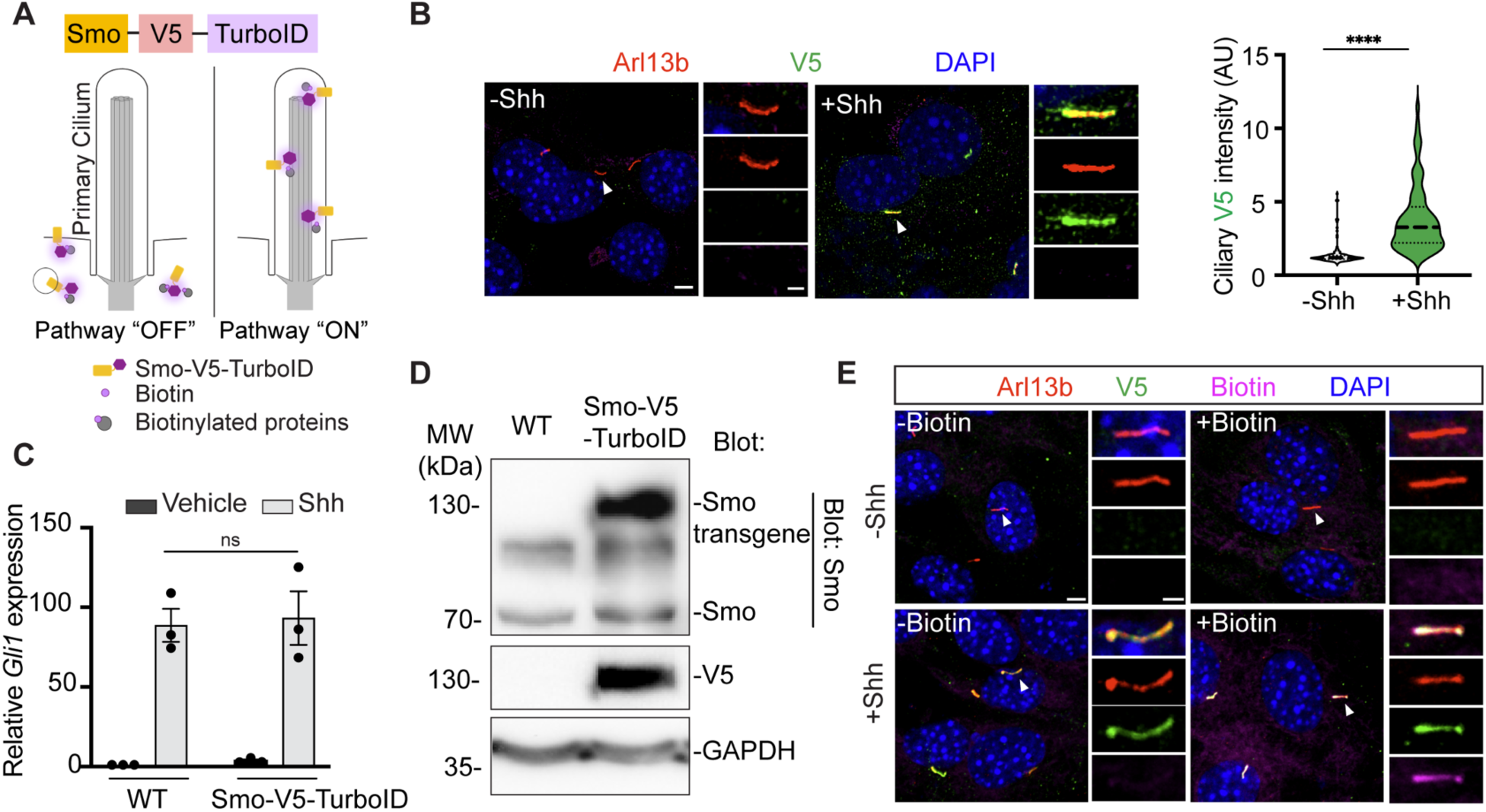
Generating the Smo-TurboID stable cell line to mimic endogenous Smo during Hh signaling activation. (A) Schematic illustration of Smo-TurboID. When the Hh pathway is off, Smo resides at the cytoplasmic membrane or other locations outside the cilium. When the pathway is on, Smo translocates to the primary cilium and is activated. TurboID biotinylates distinct cohorts of proteins at various stages of Hh activation. (B) Left: Immunofluorescence image of the selected stable cell colony showing that Smo-V5-TurboID mimics the endogenous Smo upon Shh stimulation. The transgene is highlighted by V5 staining (green); the primary cilium is marked by Arl13b (red). Scale bars, 5 μm and 2 μm (inset). Right: Quantification of ciliary V5 signal with or without Shh activation. n = 150 cells/condition; data are pooled from 3 biological replicates. Statistics: Student t-test. ****p < 0.0001. (C) Hh signaling intensity is assessed by Gli1 transcript levels in wild-type NIH3T3 (WT) and the selected Smo-V5-TurboID cell colony. No significant difference between WT and Smo-TurboID stable cell line was observed. (D) Immunoblot of WT and the selected Smo-TurboID cell colony showing that the expression levels of the transgene in comparison to the endogenous Smo. (E) Immunofluorescence image of the selected Smo-V5-TurboID cell colony after 10min biotin labeling. The biotinylated proteins are highlighted with streptavidin-Alexa 647 (magenta); V5 (green) marked the Smo-V5-TurboID transgene. Scale bar, 5 μm, 2 μm (inset).

We generated stable cell clones expressing Smo-V5-TurboID (referred to as Smo-TurboID hereafter) in NIH3T3 cells via the Flp-In system. The NIH3T3 is a cell line commonly used to study Hh signaling because it contains all vertebrate Hh signaling components. Previous studies found that overexpression of Smo in cells led to Smo self-activation and subsequent activation of Hh signaling^33,34^. To avoid this, we screened over 50 Smo-TurboID cell colonies to identify the ones that meet the following criteria. First, it shows no Smo localization in the cilium before Shh treatment (Fig. 1B). Second, it exhibits no Hh signaling activity without Shh treatment (Fig 1C). Third, the Smo-TurboID transgene expression level is comparable to the endogenous Smo (Fig. 1D). Finally, the cilium length and morphology are not affected (Fig. S1A). We selected one colony that meets all the above criteria and used this cell colony for the subsequent studies.

Then, we assessed Smo-TurboID mediated biotinylation in the selected Smo-V5-TurboID colony. Cells were treated with Shh for 16 h to induce Smo accumulation in the cilium, and labeled with biotin for 15 min before fixation for immunofluorescence staining. Salient biotinylation was detected in the cilium compared to the non-biotin condition (Fig 1E). Without Shh treatment, no Smo-TuboID is accumulated in the cilium, and no biotinylation is observed. At this resting stage, when biotin is added, Smo-TurboID at the extraciliary regions is expected to biotinylate nearby proteins, however, the signaling is too diffuse to be readily detected by immunostaining. We then purified the biotinylated proteins with magnetic streptavidin beads. Following biotin labeling, biotinylated proteins were isolated by streptavidin beads at various time point after Shh stimulation (Fig. S1B).

Together, these findings demonstrate that in the selected Smo-TurboID cell line, the transgene mimics endogenous Smo during Hh signaling activation. In addition, the expression level of Smo-TurboID enables efficient labeling of proteins in close proximity to Smo.

### Time-resolved proteomics revealed known Smo regulators

To systematically map Smo-associated proteins throughout activation of the Hh pathway, we labeled the Smo-TurboID cells at various stages of Hh activation. Previous studies suggest that Smo begins to accumulate in the primary cilia within 1 h of Shh stimulation, and reaches near-plateau levels at approximately 4 h^32^. We therefore designed a time-resolved proteomic experiment (Fig 2A). We stimulated cells with Shh for 15 min, 1 h, and 4 h, and at the end of the Shh treatment, we labeled cells with biotin for 15 min. The 15 min time point potentially captures proteins involved in the initial transport of Smo to the cilium, while the 1 h and 4 h time points allow for the identification of Smo interactors during the early and plateau phases of its ciliary accumulation. A condition with 15 min biotin but without Shh was used to capture Smo interactors at the resting stage of Hh signaling. To exclude background biotinylation, we included a non-treatment control (no biotin, no Shh) and a Shh-only control (no biotin, 4 h Shh) (Fig 2A). The enriched biotinylated proteins on streptavidin beads were digested with trypsin, followed by 6-plex isobaric tandem mass tags (TMT) labeling and mass spectrometry analysis. Three biological replicates were prepared and processed in parallel.

**Figure 2.**
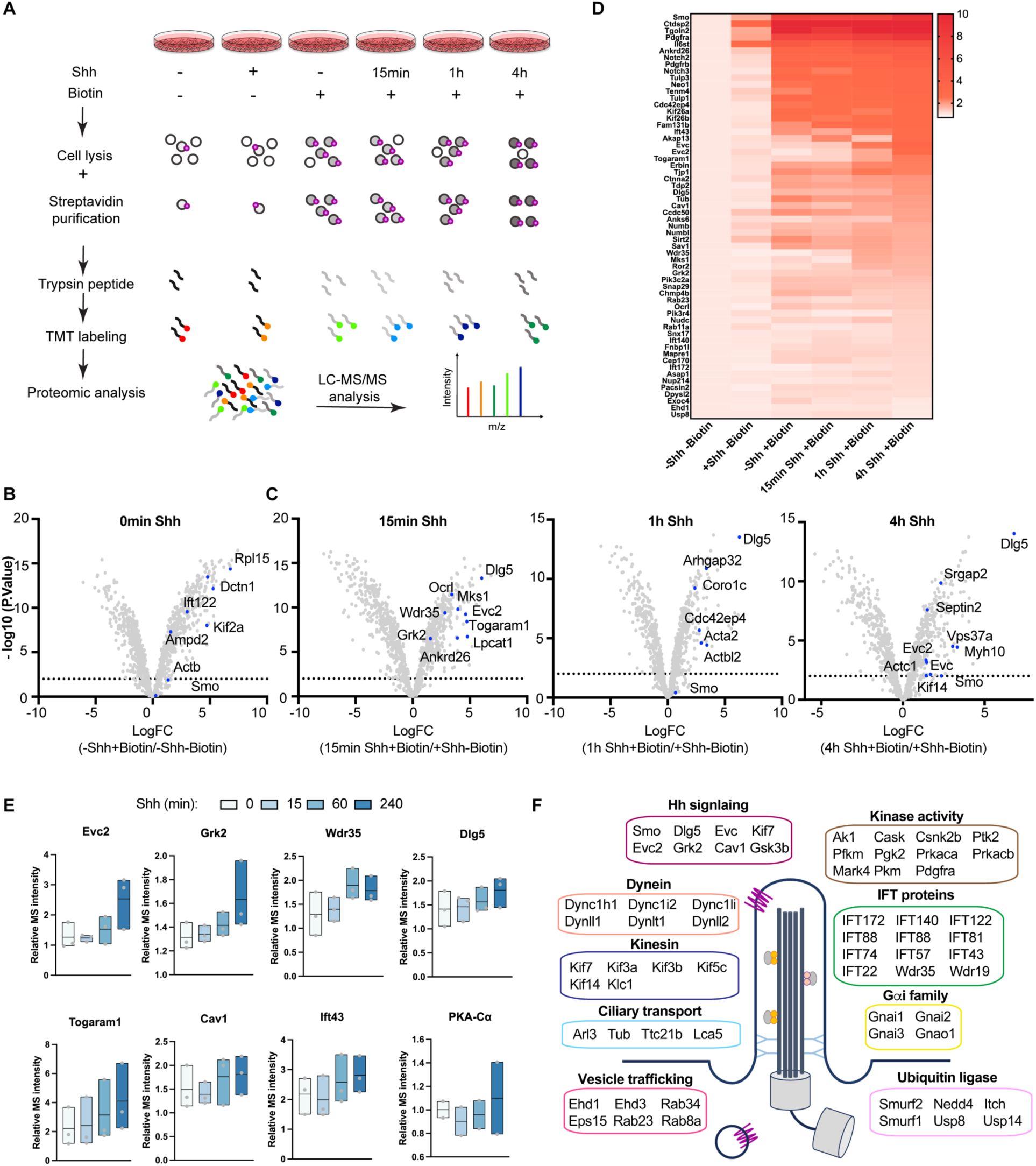
Time-resolved proteomics with Smo-TurboID revealed known Smo regulators. **(A)** Workflow of the time-resolved proteomics with Smo-V5-TurboID cell lines. Cells were grown in control conditions with biotin or Shh as indicated. For biotin labeling, cells were treated with 500 μM biotin for 15min. Shh was incubated with the cell for the specified time course. Cells were then lysed and biotinylated proteins were isolated with streptavidin beads. Biotinylated proteins were digested with trypsin on beads, and then labeled with 6-plex TMT kit. Multiplexed samples of the six channels were loaded to LC-MS/MS system for quantitative mass spectrometry. Three sets of samples, each comprising the six conditions, were prepared and processed in parallel. **(B and C)** Volcano plots showing protein fold change before Shh treatment biotin (B), and at different time points after Shh activation (C). Dash line indicates p-value 0.05. Labeled proteins are known Smo associated proteins or Hh regulators. **(D)** Heatmap of normalized protein abundance in each channel, averaged from the three sets of experiments. Top protein candidates with significant fold change and previously associated with ciliary function and Hh signaling were plotted. The color scale of relative abundances is shown on the right. (**E**) Relative mass spectrometry intensity of known regulators of Smo in Hh pathway. Mass spectrometry intensity was normalized to the control condition for each time point where no biotin was added. **(F)** Schematic view of proteins revealed in this study with known functions in the cilium and ciliary signaling.

For data analysis, we first removed common contaminants such as keratin and known endogenously biotinylated proteins from the dataset^35^. Then, we kept proteins that are detected in all six TMT channels and across all three replicates; this refined the list to 1070 proteins. Correlation of biological replicates in each TMT channel for these protein candidates demonstrated a high reproducibility across the triplicates (Fig. S2A). To normalize total protein input across the three replicates, we used an established scaling normalization method^36–38^ (see Materials and Methods), and calculated the normalized protein intensities in each replicate according to the scaling factor (Table S1). The normalized intensity was used for all the subsequent analysis.

Next, to distinguish Smo interactors at different stages of Hh signaling activation, we applied statistical analysis to compare the enrichment of proteins at each time point of Shh treatment. We analyzed the normalized protein intensities with an Empirical Bayes moderation approach, and the log-fold change (FC) was obtained with an eBayes package in R studio (Table S2, S5). Proteins exhibiting a significant increase in abundance compared to non-biotin controls (TMT ratio > 1.5) and statistically significant (P value < 0.05) are considered top candidates. Volcano plots demonstrated the enriched proteins at each time point (Fig 2 B-C). Notably, the majority of known Smo-associated proteins, as well as some Hh signaling regulators, are uncovered in our results. The relative intensities of these proteins are plotted as a heatmap (Fig. 2D, Table S3), and individual protein intensity is shown in box plots (Fig. 2E, S2B). As noted in previous studies, some proteins, such as Evc/Evc2, Grk2, Wdr35, Dlg5, Togaram1, and PKA-Cα, exhibit increased interaction with Smo following Shh stimulation^21,24–26,39–41^, while others do not show significant changes, such as Usp8 and the Gαi subunits of heterotrimeric G proteins (Gnai2)^42–45^.

Among all proteins that show higher than 1.5 TMT ratio, we identified 450 protein candidates associated with Smo before Hh pathway activation, and 576 candidates associated with Smo after Hh activation (combining all three times points after Shh treatment). The intersection of these two groups contains 112 candidates, representing proteins that bind to Smo in both conditions (Table S4). Among all candidates, 75 proteins have been reported previously to be involved in cilium function or ciliary signaling (Fig. 2F). Additional Gene ontology (GO) analysis of enriched proteins at different time points of Shh treatment highlighted diverse categories of molecular functions, including phosphatase activity before Shh treatment, and actin binding/GTPase components after Shh treatment (Fig. S2C). These results suggest that Smo interacts with distinct cohorts of proteins across various phases in Hh signaling.

Collectively, our time-resolved proteomic results with the Smo-TurboID system revealed a wide array of Smo associated proteins, offering opportunities for mechanistic studies of Smo signaling during Hh activation.

### Time-course analysis of the ciliary levels of PKA, a known Smo interactor

The PKA catalytic subunit (PKA-C) was detected in our proteomic study, with its intensity slightly increasing after Shh treatment (Fig. 2E). The interaction between Smo and PKA was originally described in the Hh pathway in *Drosophila*^14,46^, and previous cilium proteomic studies detected various PKA subunits in the cilium^47,48^. We characterized the direct interaction between Smo and PKA-C in our recent studies^16,17,19^. Here, we further determined the time course of PKA-C recruitment by Smo following the Hh pathway activation.

In the absence of Hh signaling, neither Smo nor PKA-C was detectable in the primary cilium from our immunostaining (Fig. 3A). The immunostaining signal of PKA-C in the cilium was detected 1 h after Shh stimulation, and increased significantly at the 4 h time point with the ciliary PKA-C levels coincide with the levels of ciliary Smo (Fig. 3B). Is PKA recruitment to the cilium dependent on active Hh signaling? To address this, we treated cells with cyclopamine, a small molecule Hh inhibitor that induces Smo accumulation in the cilium without being activated^49^. As expected, Smo levels in the cilium increased steadily over time after cyclopamine treatment. In contrast, no PKA-C in the cilium was detected during the entire 24 h of cyclopamine treatment (Fig. 3C-D). Thus, SMO / PKA-C colocalization in cilia requires that SMO is in an active state.

**Figure 3.**
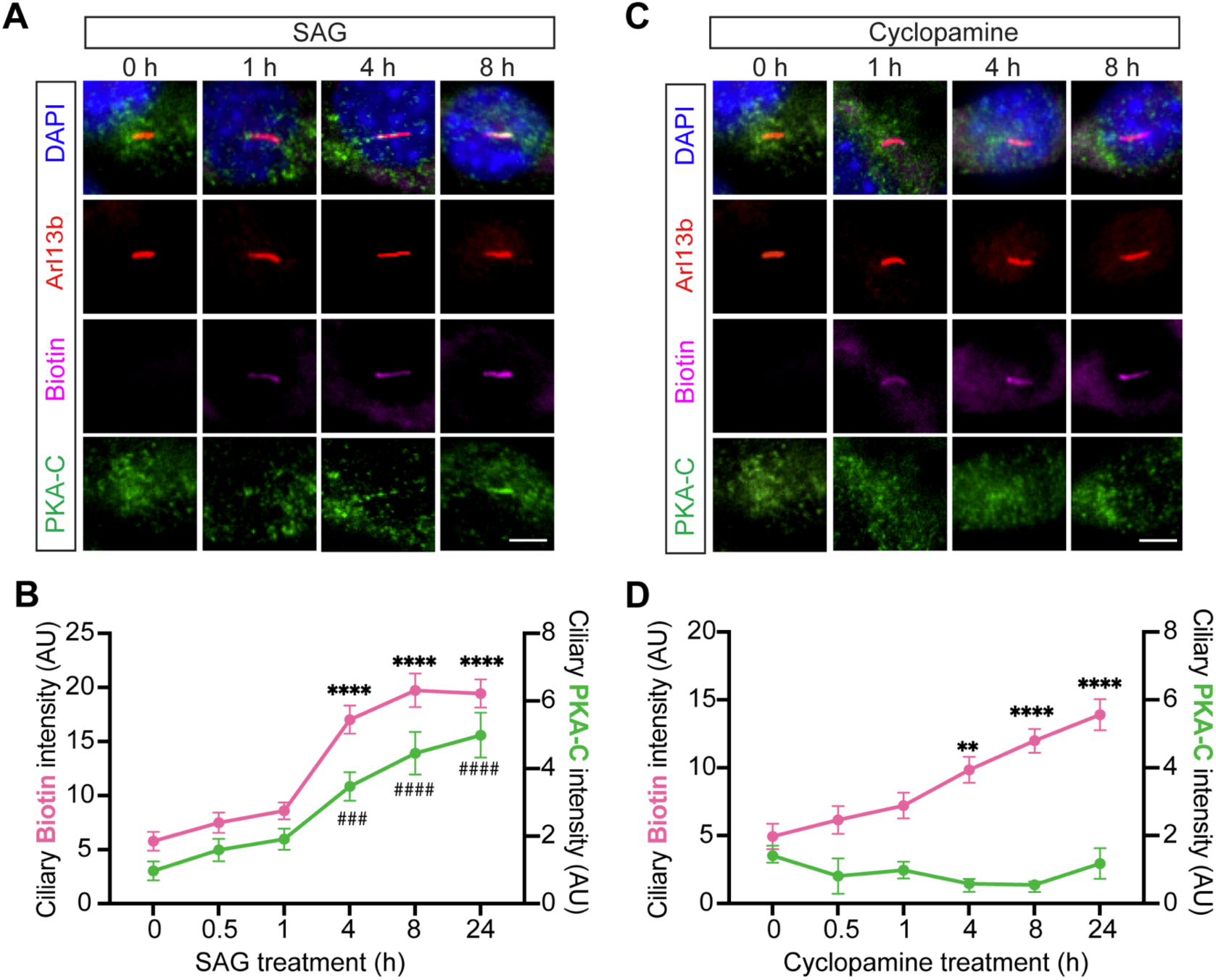
Active Smo retains PKA-C in the cilium. **(A)** Immunofluorescent imaging of endogenous PKA-C (green) in Smo-TurboID stable cell colony after 100 nM SAG treatment over the indicated time course. **(B)** Quantifications of ciliary biotin signal (pink, representing Smo-TuboID) and PKA-C signal (blue) after SAG treatment. **(C)** Immunofluorescent imaging of endogenous PKA-C (green) in Smo-TurboID stable cell colony after 5 μM cyclopamine treatment over the indicated time course. **(D)** Quantifications of ciliary biotin signal and PKA-C signal after cyclopamine treatment. n= 74 cells/condition from 3 biological replicates. Data is shown as mean ± SD. Statistics: two-way ANOVA followed by Tukey’s multiple comparison test. **p < 0.001, *** or ^###^p < 0.001, ****or ^####^ p < 0.0001, versus time 0.

To validate the specificity of the PKA-C antibody used in our study, we tested it in PKA-null MEFs in which all four PKA-C alleles (two PKA-Cα and two PKA-Cβ) are knocked out (Fig. S3A-B). When Smo-HA is expressed in wild type cells, ciliary PKA-C is detected by immunostaining. In contrast, no PKA-C signal is detected in PKA-null cells. We noted that the endogenous PKA-C in the cilium can be detected by immunostaining only in Smo-V5-TurboID cells after Shh treatment, but not in wild type parental cells (Fig. S3C-D). This is most likely due to the slightly elevated Smo levels in Smo-V5-TurboID cells compared to WT cells (Fig. 1D). This moderate increase in Smo levels does not lead to Smo self-activation, but enhances the amount of PKA-C binding to Smo after Shh stimulation. This PKA-C level in the cilium seems to be below the detection threshold in wild type cells, but is readily detectable in our Smo-V5-TurboID stable cell line.

Together, these results corroborate our findings in previous studies that Smo recruits PKA-C to the cilium in a Hh signaling-dependent manner^17,19^, and demonstrate that the Smo-TurboID system can reliably detect subtle Smo interactions that are below the detection threshold in wild-type cells.

### Git1 is a new Smo regulator that controls Smo phosphorylation status in the cilium

GO analysis highlighted actin filament binding/GTPase related proteins as top candidates of Smo interactors after Shh treatment (Fig. S2C). Among the list of new Smo interactors, we focused on one candidate, Grk-interacting protein 1 (Git1), an Arf GTPase-activating protein^50^. Git1 is shown to serves as a scaffolding protein that directs GRK2 to specific subcellular locations where Grk2 interacts with and regulates specific GPCR signaling^50,51^. Our previous study revealed a critical role of Grk2 phosphorylation on Smo at the initial stage of Hh signaling^19^, but the mechanism underlying Grk2’s translocation to the cilium remains unknown. This prompted us to investigate the potential involvement of Git1 this process.

The relative mass spectrometry intensity of Git1 in Smo-TurboID proteomic results increased after Shh stimulation (Fig. 4A). The interaction between Smo and Git1 is challenging to detect via co-immunoprecipitation (data not shown), likely due to the transient nature of this interaction. We therefore expressed YFP-Git1 in Smo-TurboID stable cell lines via lentivirus infection to facilitate the biotinylation of Git1 by TurboID. The biotinylated proteins were purified by streptavidin beads, and subsequently analyzed with western blotting. We found that Git1 is biotinylated before and after Shh stimulation but the intensity is slightly increased after Shh stimulation (Fig. 4B). This result indicates that Git1 could be in proximity to Smo in both conditions. To examine the subcellular localization of Git1, we expressed YFP-Git1 in NIH3T3 cells via lentiviral infection to ensure moderate expression level. We found that in addition to the cytosol localization, YFP-Git1 also localizes to the centrosome (Fig. 4C), which is consistent with the previously reported association with γ-tubulin^52,53^.

**Figure 4.**
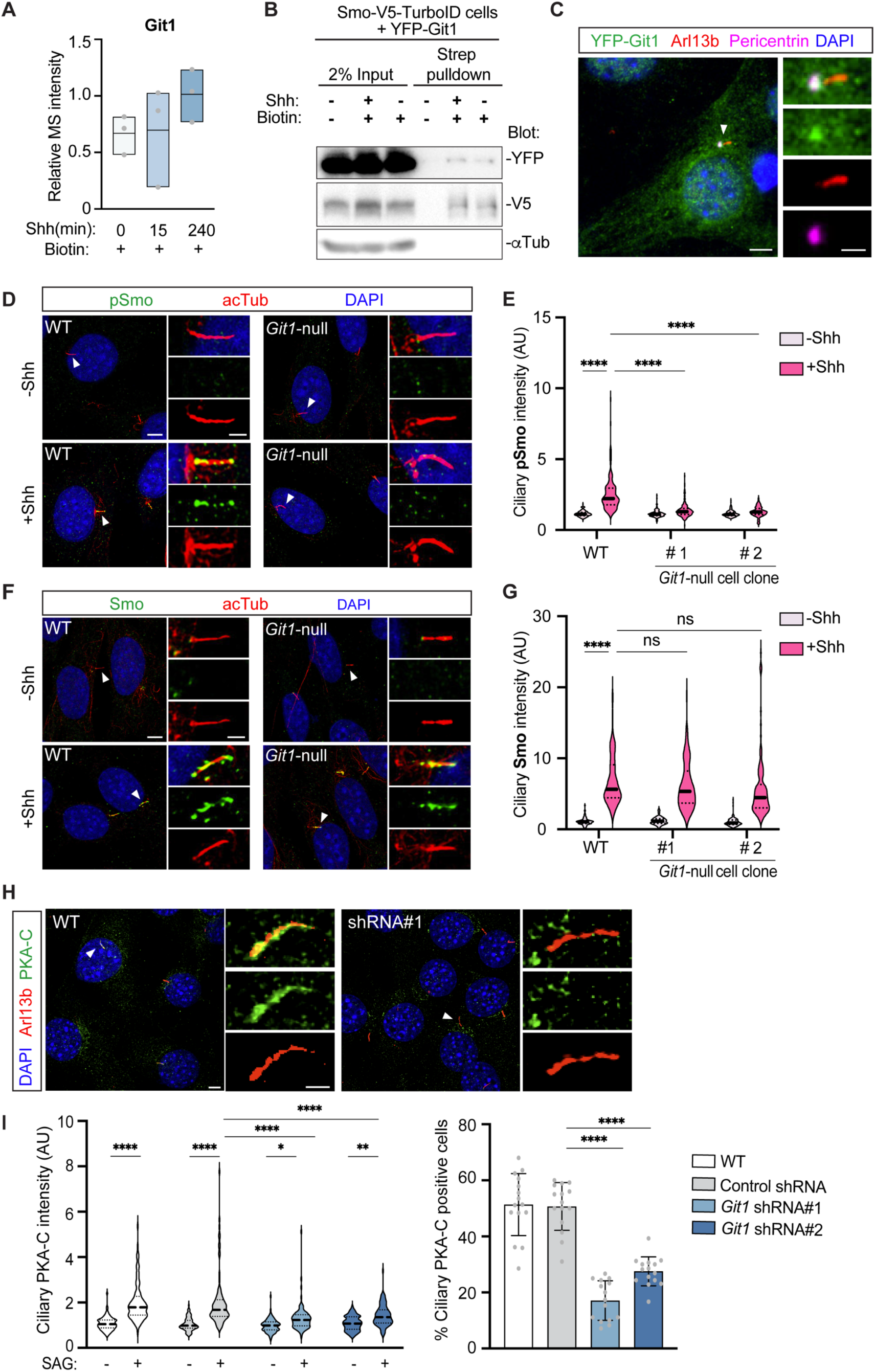
Git1 is a new Smo regulator that controls Smo phosphorylation status in the cilium. (A) Relative Git1 intensity in the Smo-TurboID proteomic results. (B) Git1 is biotinylated by Smo-TurboID independent of Shh treatment. Smo-V5-TurboID stable cells were infected with lentiviruses expressing YFP-Git1. Cells were treated with Shh for 1h, and cell lysates were used for purification with Streptavidin beads. a-Tubulin is used as loading control. (C) Git1 localizes to the basal body. NIH3T3 cells were transfected with YFP-Git1, and stained for primary cilium (Arl13b, red), basal body (Pericentrin, magenta), and nucleus (DAPI, blue). In addition to its putative distribution in the cytosol, Git1 also localizes to the basal body. Scale bar, 5 μm, 2 μm (inset). (D, F) Immunofluorescence imaging of Smo and phosphorylated Smo (pSmo) in WT and Git1-null cells. The cells are treated with 1 μg/ml recombinant Shh or vehicle. Primary cilium is highlighted by acetylated Tubulin (red). Scale bar, 5 μm, 2 μm (inset). (E, G) Quantification of ciliary Smo and pSmo immunofluorescence intensity (AU). n = 100-150 cells/condition from three biological replicates. (H) Representative images of immunofluorescence staining in Smo-TurboID cells transfected with control shRNA or shRNA against Git1. Cells were fixed 72hr after lentiviral infection and stained with PKA-C (green), Arl13b (red) and nucleus (DAPI, blue). Scale bar, 5 μm, 2 μm (inset). (I) Left: Quantification of ciliary PKA-C immunofluorescence intensity (AU), n = 90 cells/condition from three biological replicates; Right: Quantification of % ciliary PKA-C relative to total nucleus in the field. n= 15 fields per condition. Statistics in E, G, I (Left): two-way ANOVA followed by Tukey’s multiple comparison test. Statistics in I (Right) is assessed by one-way ANOVA followed by Sidak’s multiple comparison test. *p < 0.01, ***p < 0.001, ****p < 0.0001, ns, not significant.

To determine Git1’s role in Smo signaling, we generated *Git1* knockout NIH3T3 cells via CRISPR/Cas9-mediated genome editing. Sanger sequencing of the *Git1* genome sequence revealed Indel mutations that lead to frameshift or early stop codon (Fig. S4A-B), resulting in complete elimination of Git1 proteins from the two cell clones that we selected for the subsequent studies (Fig. S4C). Notably, *Git1* knockout has no impact on ciliogenesis and the cilium length (Fig. S4D-E), and no obvious defects in cell cycle progression and cell viability were found.

We then determined the impact of Git1 loss on Smo phosphorylation following Shh stimulation. Cells were stained with an antibody that specifically recognizes the Grk2 phosphorylation sites on Smo^19^. Strikingly, while Shh induced significant Smo phosphorylation in the cilia of WT cells, it failed to do so in either of the *Git1*-null cell clones (Fig. 4D-E). Next, we determined whether the reduced Smo phosphorylation is due to compromised Smo translocation to the cilium. We found that in both *Git1*-null cell clones, Shh induced comparable levels of Smo accumulation in the cilium (Fig. 4F-G). Hence, Git1 is a new Smo regulator that specifically facilitates Smo phosphorylation by Grk2 in the cilium without involving in Smo transport to the cilium.

Since PKA-C is retained in the cilium only by active Smo after its phosphorylation by Grk2, we predict that Git1 loss will impact PKA-C accumulation in the cilium. To test this prediction, we silenced Git1 expression in the Smo-TurboID stable cell line. Cells were infected with lentiviruses expressing shRNA against Git1 that effectively reduces Git1 transcript levels (Fig. S4F). 72 hr after infection, cells were stimulated with Shh and stained with PKA-C antibody. We found that cells expressing *Git1* shRNA exhibited significantly reduced PKA-C levels in the cilium compared to cells expressing control shRNA (Fig. 4H-I). Thus, Git1 loss reduced Smo phosphorylation in the cilium, which subsequently weakens Smo-PKA interaction.

### Git1 knockout reduces Grk2 transport to the primary cilium

To determine the potential interaction between Git1 and Grk2, we first examined their co-localization in the cell. We infected NIH3T3 cells with lentivirus to express Grk2-V5 and YFP-Git1. Immunostaining results demonstrated that the two proteins colocalize most prominently at the centrosome (Fig. 5A). We further confirmed the interaction of the two proteins via co-immunoprecipitation assays (Fig. 5B). These results suggest that Git1 interacts with Grk2 at the base of the cilium.

**Figure 5.**
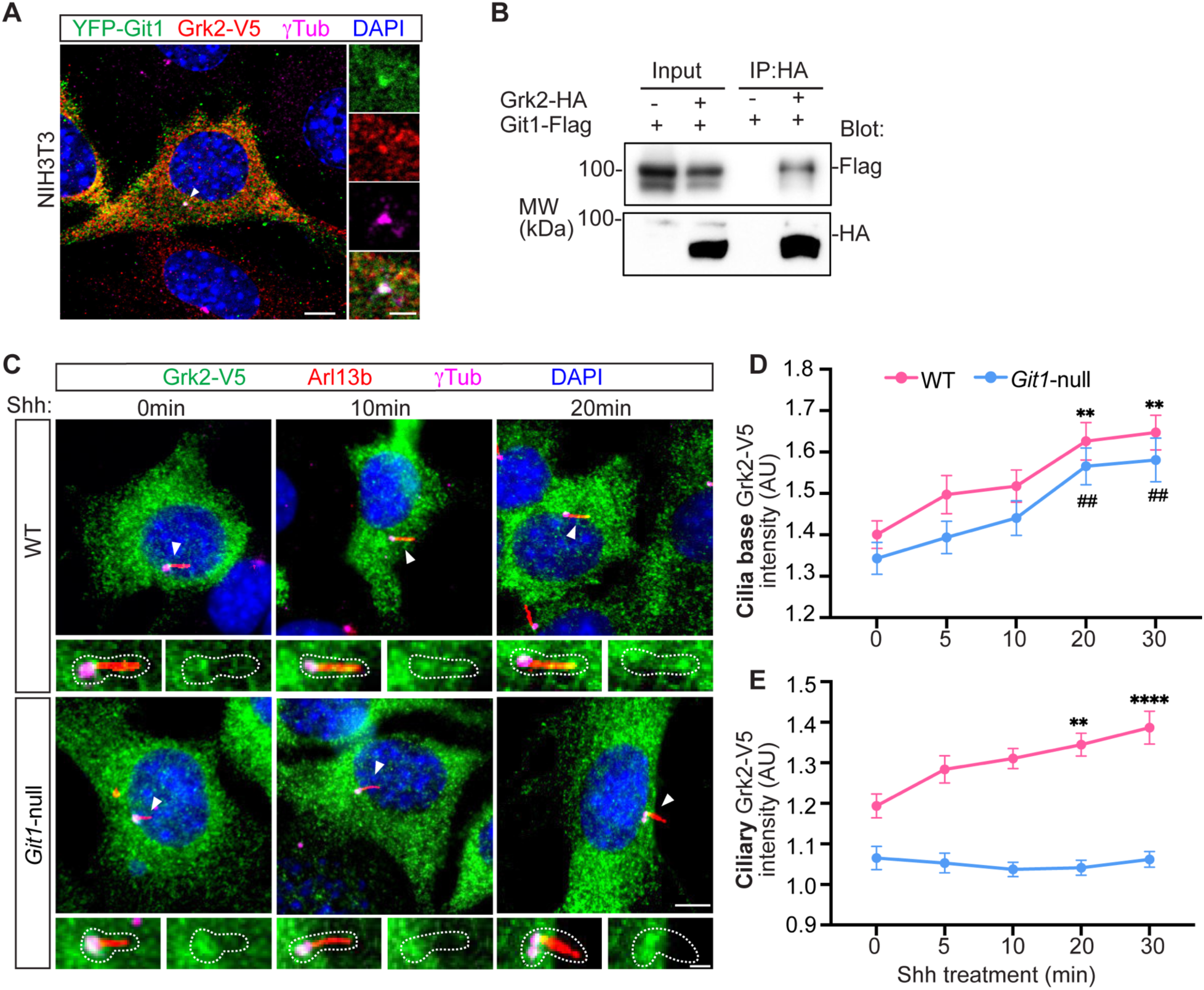
Git1 controls Grk2 transport to the cilium. **(A)** Immunofluorescence image of Grk2-V5 (red) and YFP-Git1 (green) in NIH3T3. γTub labels centrosome (magenta). Scale bars, 5 μm and 2 μm (inset). (B) Co-immunoprecipitation blot showing interaction between Git1-Flag and Grk2-V5 when co-expressed in 293T cells. (C) Representative images of Grk2 levels at the basal body and in the cilium after Shh stimulation in WT and Git1-null cells. (D, E) Quantification of basal body and ciliary Grk2 intensity in WT and Git1-null cells. n=90 cells/condition from three biological replicates. Data are shown as mean ± SD. Statistics in D and E: two-way ANOVA followed by Tukey’s multiple comparison test. ** or ^##^p < 0.001, ****p < 0.0001, versus time 0.

As reported in our previous study, Grk2 concentrates at the base of the cilium in the resting stage of Hh signaling; upon Hh signaling activation, Grk2 rapidly translocates into the primary cilium, and the ciliary Grk2 contributes to Smo phosphorylation^19^. To assess whether Git1 is involved Grk2 translocation to the cilium, we examined Grk2 localization in *Git1*-null cells. As the endogenous ciliary Grk2 level is below the detection threshold for antibody staining, we infected cells with lentivirus that expresses Grk2-V5. Lentivirus-mediated expression ensures low and uniform expression levels across the cells. We then activated Hh signaling with recombinant Shh and quantified Grk2 levels at the basal body and in the cilium shortly after Shh stimulation. We found that Grk2 levels at the basal body were slightly lower in *Git1*-null cells compared to WT cells. However, following Shh stimulation, Grk2 levels at the basal body significantly increased in both WT and *Git1*-null cells (Fig. 5C-D). In the cilium, we observed a moderate but significant 1.2-fold increase of Grk2 level in WT cells 30min after Shh stimulation. In contrast, no detectable Grk2 was observed in the cilium of *Git1*-null cells at any time point of Shh stimulation (Fig. 5C and E; note that in Fig. 5E the blue line indicates background levels of Grk2 in the cilium). It is likely that this failed Grk2 translocation to the cilium is responsible for the reduced Smo phosphorylation in the cilium (Fig. 4F-G).

Taken together, these findings suggest that Git1 is required for Grk2 entry into the primary cilium, an early event during the Hh pathway activation.

### Loss of Git1 diminishes Hh signaling

To evaluate roles of Git1 in Hh signaling, we first measured the transcript levels of *Gli1*, a Hh pathway target gene. In wild-type cells, the Smo agonist SAG induced a substantial increase in *Gli1* transcription. However, in *Git1*-null cells, SAG-induced *Gli1* transcript levels were markedly lower than WT cells (Fig 6A). At the protein level, Git1 loss also significantly reduced SAG-induced Gli1 protein expression, as detected by Western blot analysis (Fig. 6B). These results suggest that loss of Git1 diminishes Hh signaling activity.

**Figure 6.**
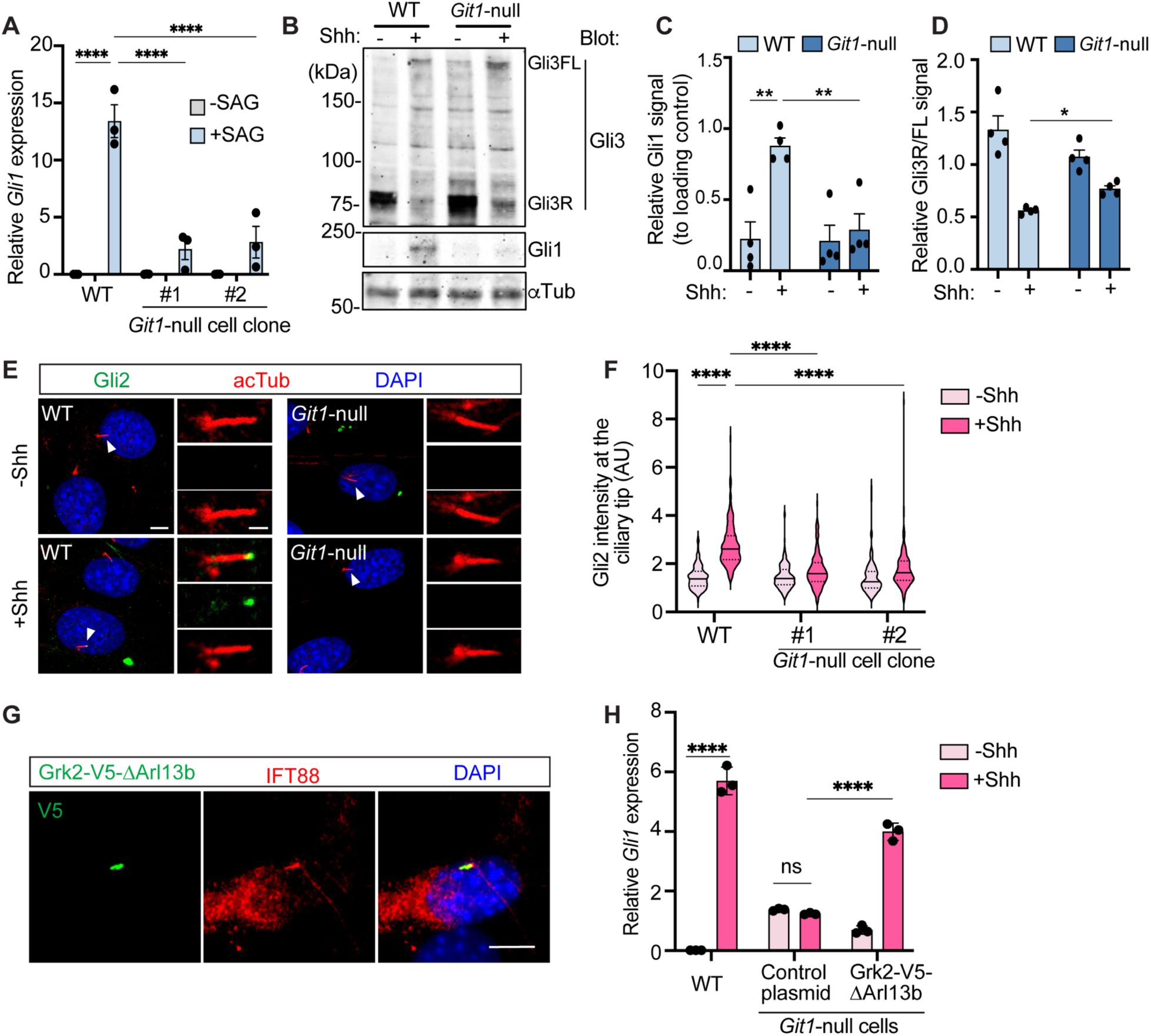
Loss of Git1 suppresses Hh signaling. (A) qPCR measurement of Gli1 transcript levels in wild-type NIH3T3 (WT) and Git1-null cell colonies. Cells were stimulated with Shh or vehicle for 24h. (B) Immunoblot of Gli3, Gli1 in WT and Git1-null cell colonies. a-Tubulin (aTub) is used as the loading control. (C and D) Quantification of immunoblot intensity of Gli3 and Gli1. n =4 independent experiments. (E) Immunofluorescent imaging of WT and Git1-null cells with or without Shh treatment. Cells were stained for Gli2 (green), and primary cilium (acTub, red). Scale bars, 5 μm and 2 μm (inset). (F) Quantification of Gli2 signal at the ciliary tip in WT and Git1-null cells. n=150 cells from 3 biological replicates. (G) Representative images of Git1-null cells infected with lentiviruses expressing Grk2-V5-Δ1Arl13b. (H) qPCR measurement of Gli1 transcript levels in WT and Git1-null cell colonies that express the indicated construct. The control plasmid refers to V5-Δ1Arl13b backbone. Statistics in A, C, D, F, H: two-way ANOVA followed by Tukey’s multiple comparison test. *p < 0.01, ***p < 0.001, ****p < 0.0001, ns, not significant.

Since Git1 loss reduces active Smo (pSmo) levels in the cilium, it may impact Gli3 processing. At resting stage, triggered by PKA phosphorylation, the full-length Gli3 is proteolytically processed into a repressor form (Gli3R) to block Hh signaling^54^. Shh stimulation leads to PKA repression, which terminates Gli3 processing and reduces Gli3R levels. We hence analyzed Gli3 processing in WT and *Git1*-null cells via Western blot. We found that in *Git1*-null cells, although Shh stimulation reduced Gli3R production, the Gli3R level is significantly higher compared to WT cells (Fig. 6B and D). Therefore, the diminished Hh signaling in *Git1*-null cells can be partially attributed to the inability to effectively suppress Gli3R production.

The activation of Gli2 is also controlled by PKA. Gli2 activation requires its transit through the ciliary tip, and loss of PKA leads to Gli2 accumulation at the ciliary tip independent of Shh stimulation^55,56^. We found that in *Git1*-null cells, Shh stimulation failed to induce Gli2 accumulation at the ciliary tip (Fig. 6E-F). Thus, reduced Gli2 activation may also contribute to the diminished Hh signaling in *Git1*-null cells.

Since loss of Git1 disrupts Grk2’s ciliary localization and the subsequent phosphorylation of Smo (Fig. 4F-G, 5C-E), we aimed to test whether targeting Grk2 to the cilium can rescue Hh signaling in *Git1*-null cells. To this end, we fused Grk2 to a cilium-targeting sequence ΔArl13b (a truncated version of Arl13b that localizes to the cilium without changing ciliary length)^23^, and expressed Grk2-V5-ΔArl13b in *Git1*-null cells via lentivirus mediated gene expression. Immunostaining confirmed localization of Grk2 in the cilium (Fig. 6G). After Shh stimulation, ciliary Grk2 significantly elevated Hh signaling intensity in Git1-null cells compared to cells expressing the control plasmids (Fig. 6H). These results suggest that Git1 regulates Hh signaling via controlling Grk2 transport into the cilium and the subsequent Smo activation.

### Loss of Git1 impairs Hh signaling in cerebellum granule neuron precursors

During cerebellar development, Hh signaling drives the proliferation of granule neuron precursors (GNPs). In the developing cerebellum, Shh is released from Purkinje neurons and acts as a mitogen to stimulate GNP proliferation^57–59^. We therefore tested roles of Git1 in Hh signaling and proliferation of primary cultured GNP.

To knockdown *Git1* in primary cultured GNPs, we infected cells with lentiviruses that express either a scrambled shRNA as control or shRNA against *Git1* (*Git1* shRNA #1, #2, Fig. 7A). Both *Git1* shRNAs efficiently reduced *Git1* expression in GNPs (Fig. 7B). To assess whether Hh signaling was affected, we stimulated cells with recombinant Shh, and measured *Gli1* transcript levels at the end of the experiment. In control shRNA-infected GNPs, Shh induced a robust *Gli1* expression; in contrast, in *Git1* knockdown GNPs, *Gli1* level was significantly reduced to only 15%-30% of control levels (Fig. 7C). Hence, *Git1* knockdown diminishes Hh signaling in primary cultured GNPs.

**Figure 7.**
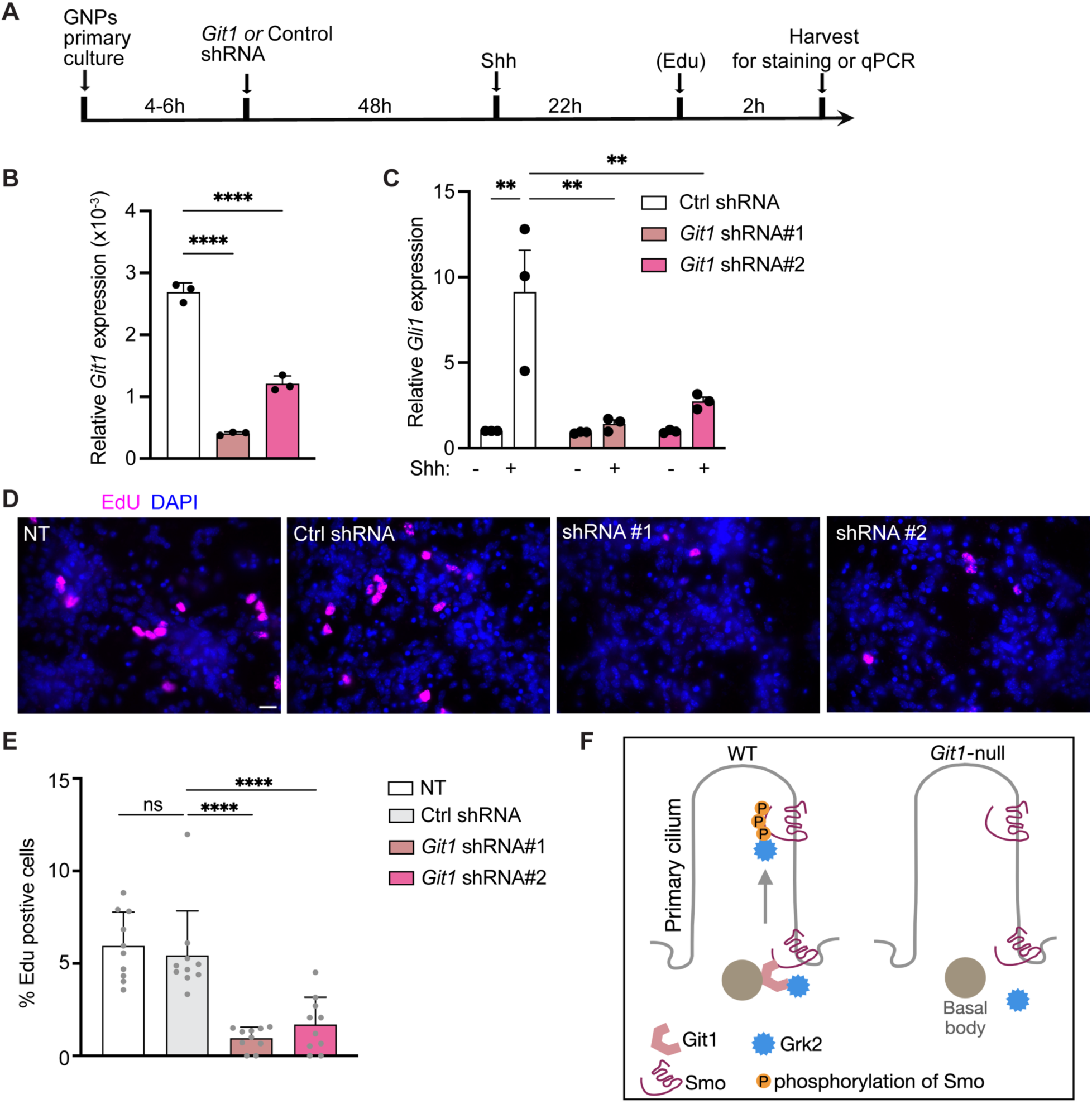
Loss of Git1 reduces Hh signaling and cell proliferation in cerebellum granule neuron precursors (GNPs). (A) Experimental workflow of GNP primary culture and treatment. (B) qPCR measurement of *Git1* transcript levels in GNPs at the end of the experiment. (C) Hh signaling intensity is assessed by Gli1 transcript levels in GNPs at the end of the experiment. (D) Immunostaining of Edu (magenta) incorporation in primary cultured GNPs. Cells are infected with lentivirus expressing control or Git1 shRNA. (E) quantification of Edu incorporation into the GNP nucleus. n = 10 fields for each condition. (F) Schematic view of Git1’s function in Hh signaling. Git1 at the ciliary base facilitates Grk2’s interaction with Smo; it also promotes Grk2 translocation into the cilium to effectively phosphorylate Smo. Without Git1, Grk2 fails to phosphorylate Smo, leading to reduced Hh signaling. Data in B, C, E are shown as Mean ± SD. Statistics: two-way ANOVA followed by Tukey’s multiple comparison test. *p < 0.01, ***p < 0.001, ****p < 0.0001, ns, not significant.

To determine the GNP proliferation rate, we performed an EdU incorporation assay. EdU was incubated with GNPs for 2 h before cells were fixed for immunostaining (Fig. 7A). We quantified the percentage of cells that incorporated EdU in randomly selected imaging fields. The results show that Git1 knockdown led to a marked reduction in EdU incorporation (Fig. 7D-E). Together, these results suggest that Git1 is essential for Hh signaling activation and cell proliferation in GNPs during cerebellar development.

## Discussion

### Smo-TurboID proteomics revealed Git1 as a new regulator of Smo phosphorylation by Grk2

Our proteomic study with Smo-TurboID identified new molecules in the vicinity of Smo during Hh signal activation. We characterized one of these new candidates, Git1, in Smo signaling. We found that without Git1, Smo accumulates in the cilium in response to Shh stimulation, similar to WT cells. However, the ciliary Smo in *Git1*-null cells is not phosphorylated by Grk2, and hence cannot effectively bind PKA-C to inhibit PKA activity (Fig. 4). Further, we found that Git1 interacts with Grk2 at the basal body; loss of Git1 abolishes ciliary translocation of Grk2 at the initial stage of Hh signaling (Fig. 5). Interestingly, in the absence of Git1, the intensity of Grk2 at the basal body does not change, suggesting that Git1 does not mediate Grk2 localization to the basal body. Taken together, our results suggest that Git1 at the ciliary base facilitates the translocation of Grk2 to the cilium at the early stage of Hh pathway activation, thereby enabling Grk2-mediated phosphorylation of Smo. Subsequently, phosphorylated Smo inhibits PKA and initiates the downstream signaling cascade (Fig. 7F). Without Git1, Smo can still translocate to the cilium, but fails to be phosphorylated by Grk2. As a result, ciliary Smo is incapable of inhibiting PKA, leading to defects in Gli3 processing and Gli2 transport at the ciliary tip.

Our results also provide important clues on the primary location where Grk2 phosphorylates Smo. Loss of Git1 does not affect Grk2 levels at the basal body, but abolishes Grk2 translocation to the cilium, and this is sufficient to block Shh-induced Smo phosphorylation in the cilium. Furthermore, targeting Grk2 specifically to the primary cilium restored Hh signaling in *Git1*-null cells (Fig. 6G-H). Together, these results strongly suggest that under physiological conditions, the ciliary shaft is the primary site where Smo is phosphorylated by Grk2. In contrast, the pool of Grk2 at the basal body is less likely contributing to Smo phosphorylation.

### An updated model of Grk2-Smo interaction facilitated by Git1

Why does the Smo phosphorylation only occur within the cilium? In other words, why does the ciliary shaft, not the basal body, serves as the primary site for Smo phosphorylation by Grk2. This could be attributed to the unique regulatory mechanisms governing Smo and Grk2 activation. With some exceptions, Grks generally recognize and phosphorylate GPCRs selectively in their active states^60^. When the Hh pathway is off, Smo remains inactive even though it transits through the cilium, because the receptor Patched prevents Smo from accessing its sterol agonists^61–63^. Only when Shh binds to Patched does the cilium environment permit Smo activation, enabling its recognition and phosphorylation by Grk2. This spatially restricted phosphorylation ensures that downstream transducers, such as PKA and Gli transcription factors, are properly regulated within the cilium to reliably propagate the signaling cascade.

In our previous study, we observed that when inactive Smo accumulates in the cilium by cyclopamine treatment, SAG can induce significant Smo phosphorylation in the cilium within 20 min^19^. These results suggest that Grk2 rapidly translocates into the cilium following Hh pathway activation to phosphorylate Smo. In the current study, we found that Git1 interacts with both Smo and Grk2 at the cilium base, and is required for Grk2 translocation to the cilium. Taken together, we propose an updated model of Grk2-Smo interaction. Git1, a scaffold protein at the basal body, transiently binds to both Smo and Grk2 to bring the two molecules together, a process that could be enhanced by Hh signaling activation (Fig. 4A-B). However, as Smo is only activated within the cilium, Grk2 needs to enter the cilium to recognize and phosphorylate active Smo (Fig. 7F). As Grks typically bind directly to their GPCR substrate without the need of intermediary molecules, Git1 does not enter the cilium together with Smo and Grk2.

As soluble proteins, Grks require specific mechanisms to bring them to the vicinity of GPCRs. Some Grks, such as GRK1, associate with membrane via the prenylation modification at the C-terminus^64^. GRK2/3 has been shown to be recruited by Gβγ to the membrane to gain access to their GPCR substrates^65–67^. However, evidence of Gβγ in the vicinity of Smo at the ciliary base is lacking, and our previous study showed that of Gβγ is dispensable for Smo phosphorylation in the cilium^19^. Anchored to the basal body by γ-tubulin^53^, Git1 is located at an ideal position to serve as a scaffold to bring Grk2 close to Smo.

### Git1 acts as a positive regulator of Hh signaling

Git1 may regulate Hh signaling at locations beyond the cilium base. Git1 is a multi-domain containing protein, and previous studies implicate Git1 in a variety of cellular processes^50^. Its C-terminal coiled-coil and paxillin-binding domains are known to mediate protein-protein interactions; its N-terminal ArfGAP domain has been shown to regulate GPCR internalization and trafficking^27^. The membrane-associated Git1 has been shown to play important roles in Grk2-mediated receptor endocytosis and recycling^27,51^. The Smo-associated Git1 outside the cilium may function to prevent premature internalization of Smo from the cytoplasmic membrane. As Smo translocates to the cilium via lateral movement from the adjacent plasma membrane^68^, Git1 might help maintain this pool of Smo at the plasma membrane so that Smo is available for ciliary localization once the pathway is on. A similar mechanism of regulating the plasma membrane levels of Smo has been reported as the action mechanism of the ubiquitin system of MEGF8, MGRN1^69^.

Our findings in NIH3T3 cells are corroborated by previous studies on *Git1*-null animal models. Three *Git1*-null mouse models have been reported^29,30,70^, and all mouse models exhibited 50-60% postnatal lethality. The surviving mice exhibited abnormality in multiple tissues and organs, including microcephaly, reduced pulmonary blood vessels, altered cortical layering, and cerebellar agenesis^29^. These phenotypes are consistent with diminished Hh signaling in these tissues. The *in vivo* phenotypes are less pronounced compared to the reduced Hh signaling in Git1-null NIH3T3 cells, suggesting that in mouse models, the Smo-Grk2 interaction may also be facilitated by proteins with similar functions as Git1. Alternatively, Git1 could be required for Smo-Grk2 interaction in some tissues but not others.

### The time-resolved proteomic dataset revealed Smo regulators over the course of Hh signaling activation

Our time-resolved proteomic study with TurboID identified distinct categories of proteins in the vicinity of Smo at different stages of Hh signal activation. These results provide a valuable resource for investigating the signaling mechanisms of Smo, as demonstrated by our characterization of Git1’s role in Smo phosphorylation. Compared to conventional affinity purification, proximity labeling-based proteomics has the strength of capturing transient and weak interactions. It is worth noting that our proteomic results recapitulated nearly all known Smo-interacting proteins, and also uncovered new Smo-associated molecules. The Smo-TurboID system detected Grk2 association with Smo at 15 minutes of Shh stimulation, consistent with the previous finding using optical imaging approaches, which identified this interaction as an early event in Hh activation^19^. Interestingly, the most abundant proteins at early time points (15 min and 1 h after Shh stimulation) are actin binding, GTPase binding, and cytoskeleton motor proteins (Fig. S2), suggesting the involvement of protein transporting machinery during these stages. At later stages, enriched molecules are proline-rich domain containing or SH3-domain containing proteins, and phosphatidylinositol binding proteins. Investigating how these distinct groups of Smo-associated proteins function in different stages of Hh transduction represents an important direction for future research.

**Figure S1.**
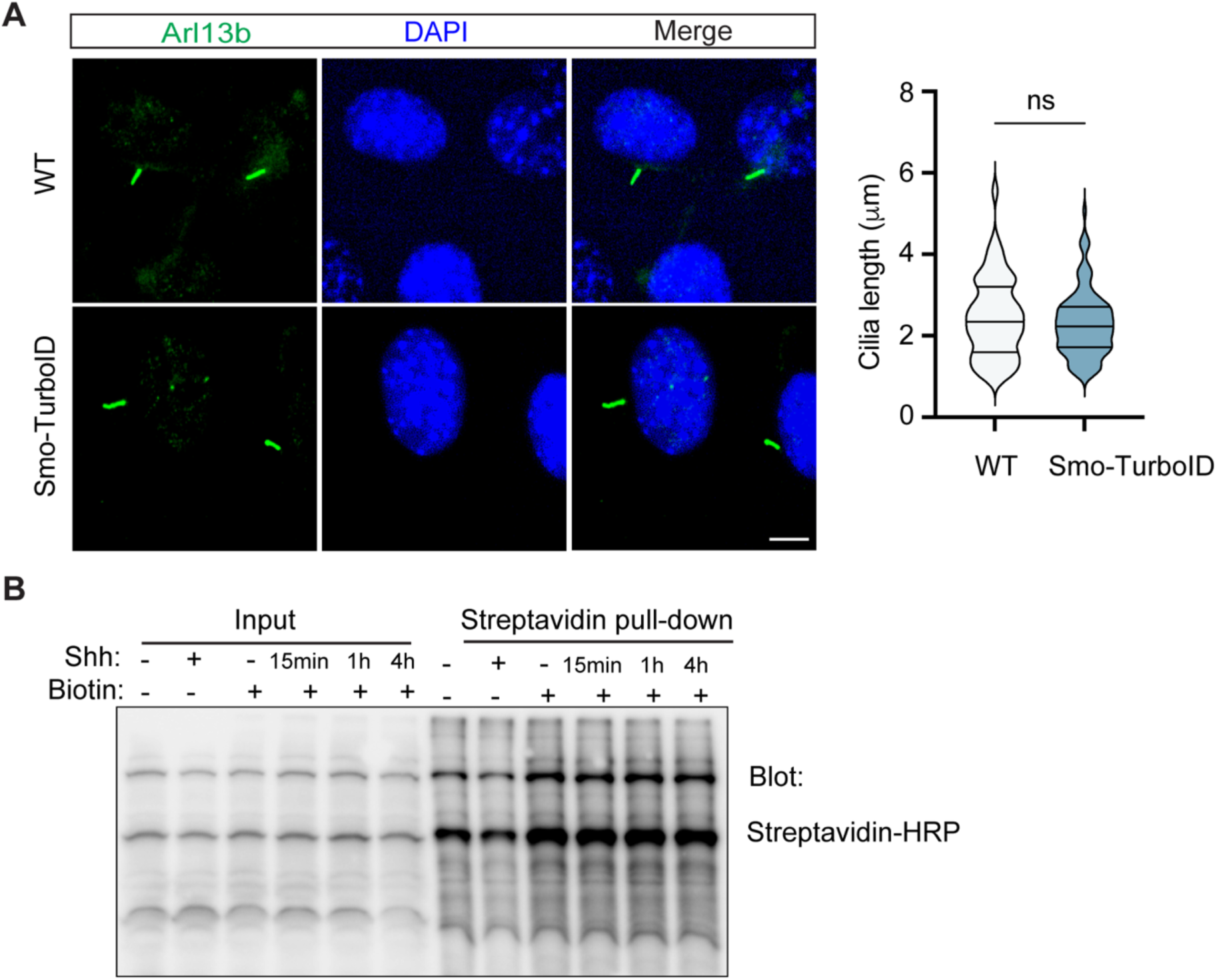
Generating the Smo-TurboID stable cell line to mimic endogenous Smo during Hh signaling activation. (A) Left: Staining of the primary cilium in WT and the selected Smo-V5-TurboID cell colony. The cilia are highlighted by Arl13b (green). Scale bar, 5 μm; Right: Quantification of the cilium length. Statistics: Student’s t-test. ns, not significant. (B) Purification of biotinylated proteins from the Smo-TurboID cells. The selected Smo-V5-TurboID cell clone was treated with Shh for the indicated duration and labeled with biotin for 10 minutes. Cell lysates were then subjected to purification using streptavidin beads, followed by Western blot analysis.

**Figure S2.**
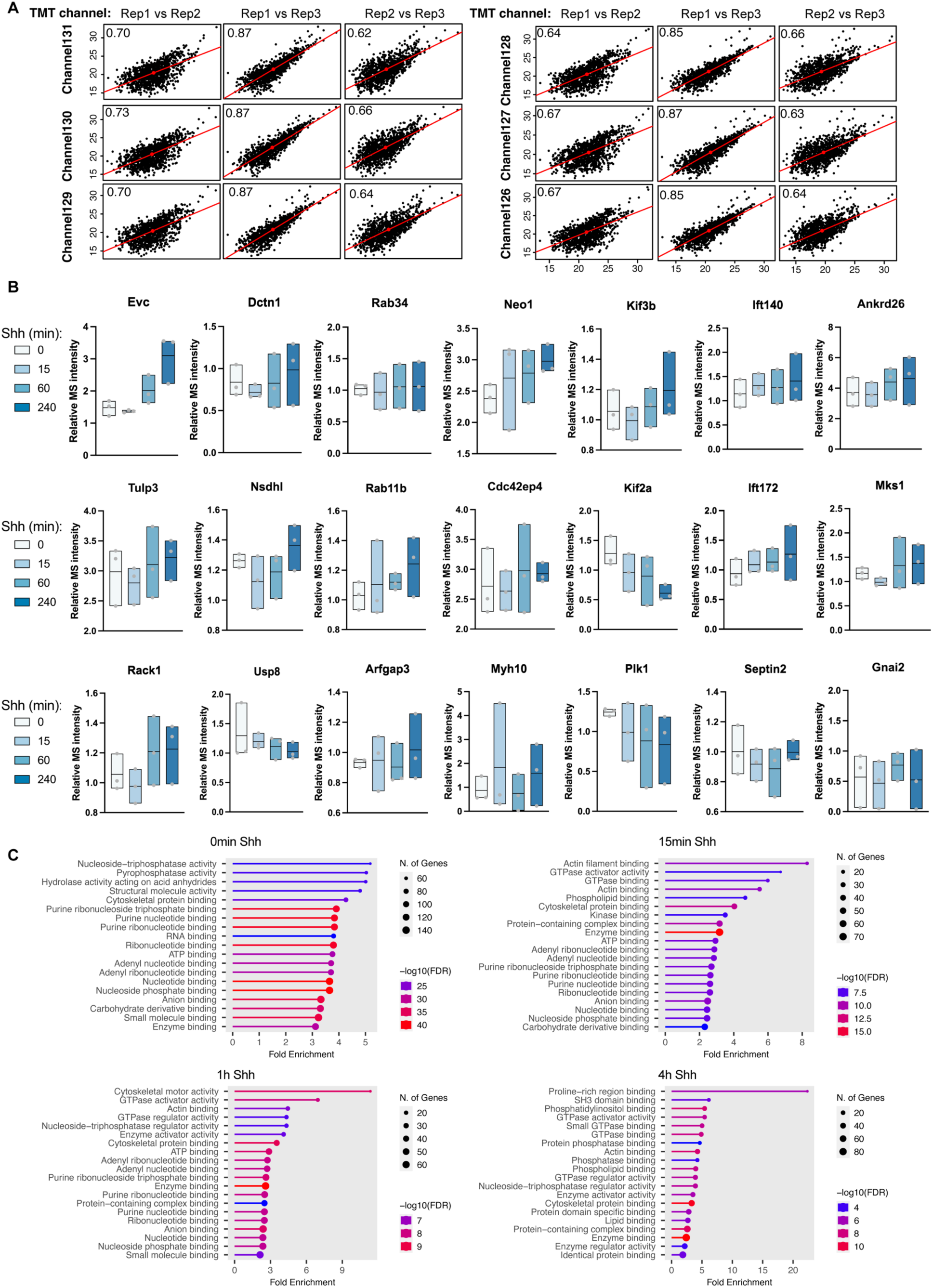
The overview of time-resolved proteomics with Smo-V5-TurboID stable cell line. (A) Correlation of biological replicates for the six conditions: channel 131 (+Shh 4 h, +Biotin), channel 130 (+Shh 1h, +Biotin), channel 129 (+Shh 15 min, +Biotin), channel 128 (-Shh, -Biotin), channel127 (-Shh, +Biotin), channel126 (+Shh, - Biotin). 1070 proteins that were detected in all six channels across 3 replicates are plotted. (B) Mass spectrometry intensity of known Hh signaling regulators revealed in this study. Mass spectrometry intensity was normalized to the control condition for each time point where no biotin was added. (C) Gene ontology analysis of molecular function for proteins with mass spectrometry ratio > 2.

**Figure S3.**
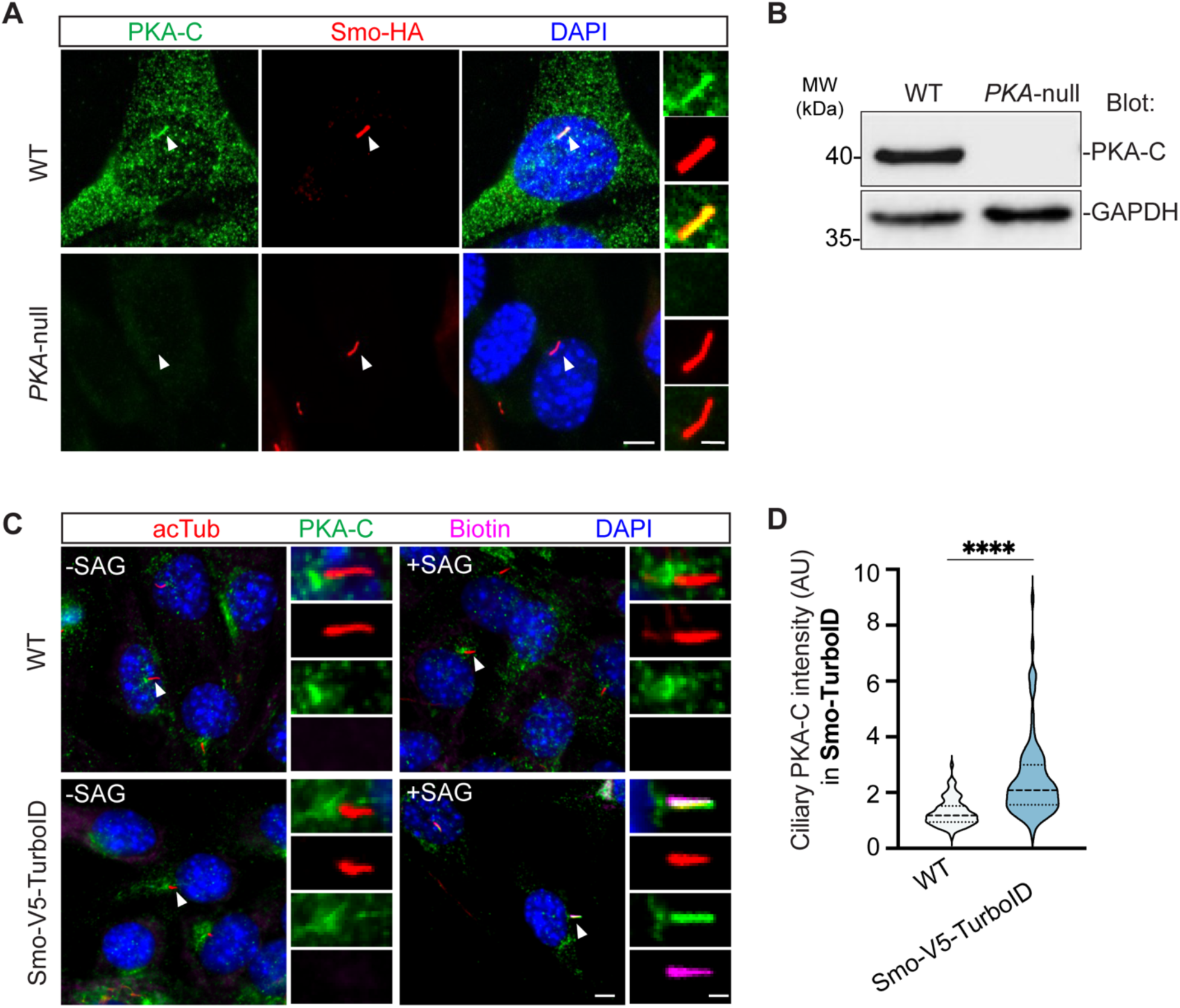
Overexpression of Smo recruits PKA-C to the primary cilium. **(A)** WT or PKA-null MEF cells were transfected with Smo-HA, treated with Shh for 16hr, and stained for HA (red), PKA-C (green) and DAPI (blue). Overexpressing Smo recruits PKA-C to the primary cilium in WT cells, but not in PKA-null MEF cells, indicating the specificity of the PKA-C antibody in immunofluorescence staining. **(B)** Immunoblot for PKA-C in WT and PKA-null MEF cells. GAPDH was used as loading control. **(C)** Immunofluorescent imaging of endogenous PKA-C (green) in wild-type and Smo-V5-TurboID stable cell line with or without 100 nM SAG treatment for 24h. Cilia is marked by acetylated-tubulin (acTub, red); Smo-TurboID intensity is indicated by biotin labeling (magenta); DAPI (blue) marks nucleus. **(D)** Quantification of PKA-C intensity in the primary cilium of WT and Smo-V5-TurboID stable cell line. n= 90 cells from 3 independent experiments. Statistics: Student t-test.

**Fig. S4.**
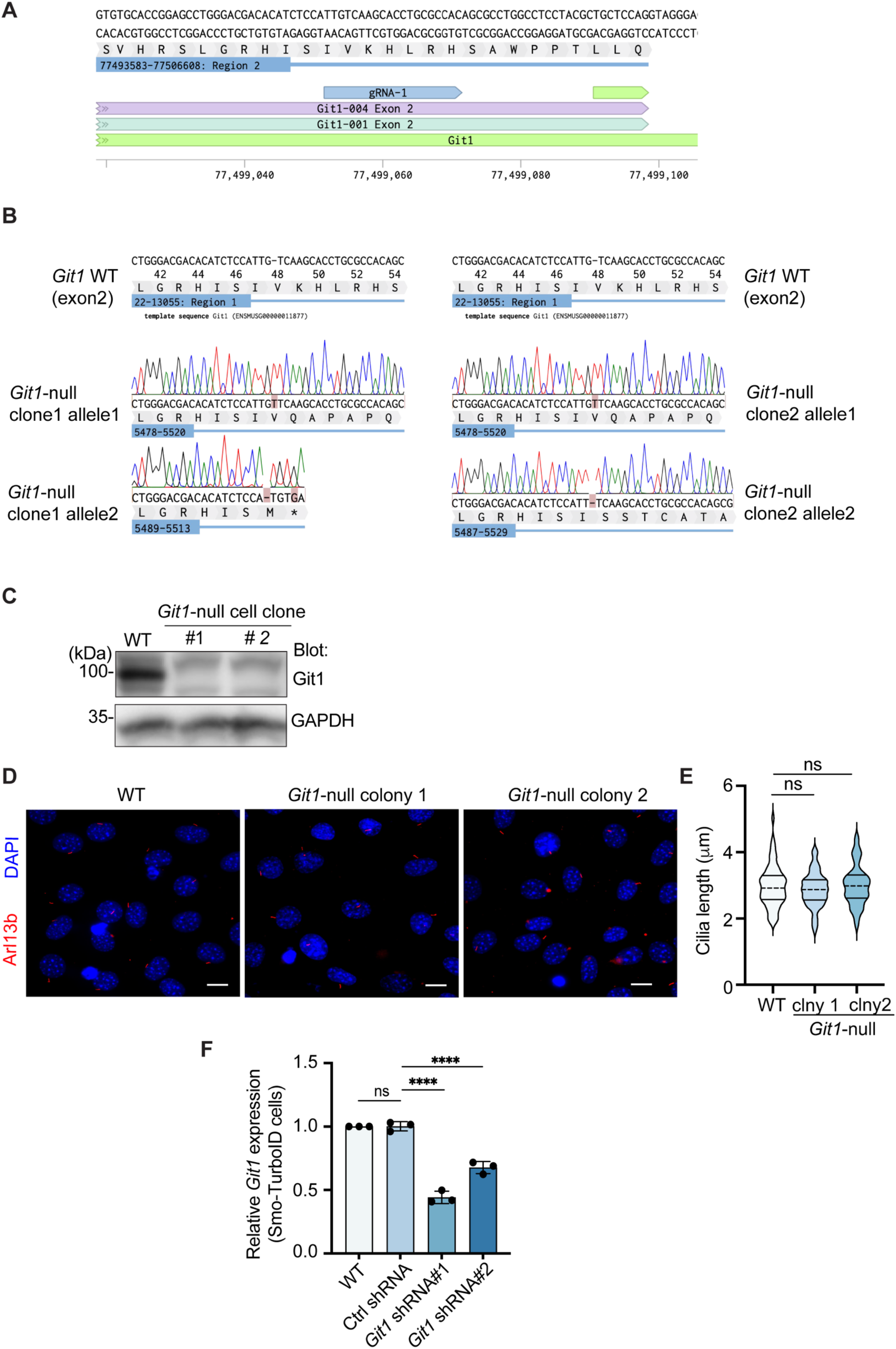
CRISPR/Cas9 mediated Git1 knockout in NIH3T3 cells. (A) guide RNA was designed to target exon 2 of mouse *Git1*. (B) The gRNA targeting region in mouse genomic was amplified by genomic PCR, ligated into TOPO vector, and transfected into chemically competent cells. 20 bacterial colonies of each cell clones were randomly picked and sequenced. The Sanger sequencing results were aligned with the genome sequence of the M. musculus. Single base pair deletion and insertion are identified, resulting in biallelic frameshift or early termination. (C) Immunoblot of Git1-null cell lines showing that Git1 protein is not detected in knockout cells. GAPDH is used as the loading control. (D) Cilium staining in WT and Git1-null cell colonies. Primary cilium is marked with Arl13b (red). Scale bar, 10 μm. (E) Quantification of cilium length, n = 60 cells/condition from 3 biological replicates. (F) Git1 transcript levels in cells transfected with shRNA against Git1. Statistics in (E-F): one-way ANOVA followed by Sidak’s multiple comparisons test. *p < 0.01, ***p < 0.001, ****p < 0.0001, ns, not significant.

## ACKNOWLEDGEMENTS

We thank Dr. Ljiljana Milenkovic from Matthew Scott lab for the antibodies against Smo. TurboID is provided by Dr. Alice Ting’s lab (Stanford University). We thank Dr. David Gravano (Stem Cell Instrumentation Foundry, University of California, Merced) for fluorescence-activated cell sorting. The *PKA-C*α knockout MEF cell is a gifted from the Anderson lab at Sloan Kettering Institute, and was used to generate the double knockout in PKA-Cα and Cβ. Research in the laboratory of X.G. was supported by NIH/NCI R15CA235749, NIH/NCI R21CA274595, NIH/NIGMS R01GM143276, and NSF CAREER award IOS-2143711. This project is supported by NIH/NIGMS P41GM103533 to J.R.Y., and NIH/NIGMS R35GM133672 to B.R.M.

## AUTHOR CONTRIBUTIONS

X. Ge conceived the project. X. Ge and J. Zhang designed the experiments. J. Zhang performed most experiments, analyzed, and interpreted the data. G. Kaur contributed to the Co-IP experiments and characterization of protein-protein interaction. E. Cai participated in data analysis and figure construction. O. T. Gutierrez contributed to the analysis of cilium protein quantification. X. Liu participated in the preparation of mass spectrometry samples and data analysis. S. Baboo and JK Diedrich processed samples for mass spectrometry and performed the initial analysis of the mass spectrometry results, under the supervision of J.R. Yates. J-F Zhu generated PKA double knockout cells under the supervision of B. R. Myers. X. Ge, J. Zhang and B. R. Myers wrote the manuscript. All authors revised the manuscript.

## MATERIALS AND METHODS

### Antibodies

The following antibodies were used in this study: Rabbit anti-pSmo (7TM antibodies, 7TM0239A-IC), Rabbit anti-Smo (gift from M. Scotts, Stanford University), Mouse anti-acetylated tubulin (Sigma,T6793), Rabbit anti-Arl13b (Proteintech, 17711–1-AP), Rat anti-Arl13b (BiCell Scientific, 90413), rabbit anti-IFT88 (Proteintech, 13967-1-AP), Goat anti-Gli2 (R&D Systems, AF3635,), Goat anti-Gli1 (R&D Systems, AF3455), Rabbit anti-Git1 (Novus Biologicals, NBP1-86144), Chicken anti-GFP (Aves labs, GFP-1020), Rabbit anti-GFP (Thermo Fisher Scientific, A-11122), Mouse-anti-Flag (Sigma, F3165), Rabbit anti-HA (Cell Signaling Technology, 3724), Mouse anti-GAPDH (Thermo Fisher Scientific, MA5-15738), Mouse anti-V5 (Thermo Fisher Scientific, R960-25), Mouse anti-PKACα (BD Biosciences, 610980), Rabbit anti-PKACα ( Cell Signaling, D38C6), Mouse-anti-pericentrin (BD Biosciences, 611814), Mouse anti-gamma Tubulin (Proteintech, 66320-1-Ig), DAPI ((Thermo Fisher Scientific, D21490), Donkey anti-rabbit Rhodamine (Jackson ImmunoResearch Labs, 711-025-152), Donkey anti-rabbit Alexa 488 (Jackson ImmunoResearch Labs, 711-545-152), Donkey anti-mouse Alexa 488 (Jackson ImmunoResearch Labs, 715-025-151), Donkey anti-mouse Rhodamine (Jackson ImmunoResearch Labs, 715-545-151), Donkey anti-Chicken AlexaFluor 488 (Jackson ImmunoResearch Labs, 703-545-155), Donkey anti-Goat AlexaFluor 488 (Jackson ImmunoResearch Labs, 705-545-003), Donkey anti-Goat AlexaFluro 647 (Jackson ImmunoResearch Labs, 705-605-147), Alexa Fluor 647 Streptavidin (Jackson ImmunoResearch Labs, 016-600-084), Donkey-anti, HRP-Conjugated Streptavidin (Thermo Fisher Scientific, N100).

### Cell line generation, cultivation and manipulation

PKA-deficient MEFs were obtained from Kathryn Anderson’s lab^55^ and determined to be PRKACA^+/-^; PRKACB^−/-^ based on Western blotting with PKA-Cα and PKA-Cβ specific antibodies (a gift from Mark Knepper, NIH). The remaining PRKACA allele was then knocked out using the CRISPR-mediated gene disruption technique (Alt-R system, IDT) as previously described^16^. Briefly, RNP complexes using Alt-R predesigned CRISPR/Cas9 cRNA against PRKACA (/AltR1/rUrC rUrCrC rCrArC rCrUrA rCrGrG rCrGrG rArUrG rUrUrU rUrArG rArGrC rUrArU rGrCrU /AltR2/) were delivered to PKA-deficient MEFs via a Neon Electroporation system (Thermo Fisher). Two days later, cells that had taken up the RNP (identified as Atto550+ cells, via the Atto550-labeled crRNA in the RNP complex) were single-cell sorted by FACS, expanded, and tested for loss of PRKACA via Western blotting. Flp-In 3T3 cells ((Thermo Fisher Scientific, R76107) and 293 T cells (ATCC, CRL-3216) were cultured in DMEM (supplemented with 10% Fetal bovine serum) according to manufacturer’s instructions. PKA-Ca null MEFs were cultured in DMEM (supplemented with 10% Fetal bovin serum). NIH3T3 cells (ATCC, CRL-1658) were cultured in DMEM (supplemented with 10% calf serum) as previously described. Ciliation was induced by reducing the growth media to 0.5% serum for 16-24h. To induce Hh signaling, growth media were supplemented with 100 nM SAG, 1 µg/ml recombinant ShhN or ShhN condition medium (20%-30% [vol/vol]) depending on batch) produced with 293 ecR-Shh-N cells (gift from R. Rohatgi, Stanford University). To block Hh signaling, 5 µM cyclopamine (Selleckchem, S1146). Transfection were performed using Lipofectamine 2000 (Invitrogen) according to manufacturer’s instructions. *Git1* gene was disrupted in NIH3T3 cells using CRISPR/Cas9-mediated genome editing targeting exon 2. Clones of each cell line were obtained by single-cell sorting. Clones with disrupted gene were screen with immunofluorescence and western blotting using protein-specific antibodies. Selected cell clones were characterized by sequencing to confirm missense mutation leading to early termination of translation and frame shift mutation. Flp-In 3T3 cells stably expressing Smo-V5-TurboID were generated using Lipofectamine 2000 transfection.

### DNA constructs

For generation of Smo-V5-TurboID cell line, full-length mouse Smo was first cloned into pEF5/FRT/V5-DEST backbone (Thermo Fisher Scientific, V602020). Then TurboID (gift from A. Ting, Stanford University) was attached to the C terminus of Smo and linked by V5 tag. To observe localization and activity of Git1, YFP was fused to the N terminus of human Git1 (Addgene, 15225) and cloned into FUGW backbone (Addgene, 14883). To observe Grk2 activity, bovine Grk2 (gift from B. Myers, University of Utah) was cloned into FUGW backbone and linked by a V5 or HA tag. To target bGrk2 to the primary cilium, a truncated version of Arl13b (ΔArl13b) described previously in Liu et al. 2024 was attached to the C terminus of Grk2 and linked by V5 tag.

### Primary culture of cerebellum granule cell precursors (CGNPs)

Cerebelluar GNPs were cultured as previously described^71^. Briefly, cerebella from postnatal day 7 (P7) C57BL/6J mice were cut into small pieces and incubated at 37°C for 20min in 15U/ml papain solution (Worthington Biochemical Corporation, LS003126) and DNase I (Roche, 11284932001) in Hanks’ Buffer with 20 mM Hepes (HHBS). HHBS was used to rinse tissue once and then removed. Tissues were then triturated in Neurobasal medium (Gibco, Cat# 21103049) containing DNase I to obtain single cell suspension. Cells were centrifuged at 1000 rpm for 5 min at 4°C and resuspended in Neurobasal medium containing B-27 Supplement (Gibco, 17504044), GlutaMAX Supplement (Gibco, 35050061) and 1% Pen Strep (Gibco, 15140122). Cells were plated on Poly-D-Lysine containing Laminin (Gibco, 23017015) coverslips at 1.2 × 106 cell/ml. After 24h in culture, cells were treated with DMSO or 100 nM SAG for 24h and fixed in 4% PFA for immunostaining. GNPs were blocked with blocking buffer (0.2% Triton X-100, 2% Donkey serum in PBS) for 1 h at room temperature. After blocking, cells were incubated with rabbit anti-pSMO (1:1000), rat anti-ARL13B (1:500) at 4°C overnight. Subsequently, cells were incubated with secondary (ThermoFisher Scientific) for 1 h and Hoechst 33342 for 10min at room temperature. Cells were mounted in Fluoromount-G. Imaging of GNPs was done using YOKOGAWA CSU-W1 system with PHOTOMETRICS PRIME 95B camera with 100X oil immersion lens.

### TurboID labeling experiments

Cells were incubated in the presence of 500 μM biotin diluted in DMEM for 15min before harvesting. For non-labeling conditions, pure DMEM were added. After biotin labeling, the medium was aspirated quickly and washed three times with ice-cold DPBS and lysed in subcellular fractionation buffer (20 mM HEPES, 10 mM KCl, 2 mM MgCl_2_, 1mM EDTA, 1 mM EGTA, 1mM DTT, pH 7.5 and protease inhibitors). Cells were then passed through 27-G needles and centrifuged at 720 x g (3000 rpm) for 5 min to separate nuclei and the supernatant were collected. Supernatant were respun at 10,000 x g (8000 rpm) for 5 min and collected to further remove debris and nuclei. Samples were then supplemented with RIPA including 0.5% NP-40, 0.1% SDS, 0.5 % sodium deoxycholate and protease inhibitors was added to further lyse the cells. cells were then sonicated and centrifuged at 10,000 x g for 10 min and supernatant was collected.

### Streptavidin purification

Protein concentration from each condition were normalized to equal concentration and volume using BCA assay. Samples were added to washed and equilibrated streptavidin magnetic beads (Thermo Fisher Scientific, 88816) and incubated for 1.5 hr at room temperature. Unbound material was removed, and samples were proceeded with western blot (WB) analysis. Beads with bound proteins were washed extensively with a series of buffer to remove nonspecific binders. Briefly, samples on beads were washed with RIPA twice, 1 M KCl once, 0.1 M Na_2_CO_3_ once, 2 M Urea (in 10 mM Tris-HCl, pH 8.0) once, RIPA twice and DPBS once. For western blot, beads were eluted with 2 x SDS sample buffer.

### On-bead trypsin digestion of biotinylated proteins and TMT labeling

Proteins bound to the magnetic beads were denatured with 8 M urea, reduced with tris (2-carboxyethyl) phosphine (TCEP), alkylated with 2-chloroacetamide, and precipitated with methanol-chloroform^72^. Bead-bound proteins were digested with trypsin and the peptides labeled with TMT 6-plex (Thermo). TMT-labeled peptides were pooled and fractionated into 8 fractions at high pH (Pierce, 84868) according to the manufacturer’s instructions.

### Mass spectrometry

Tryptic peptides were separated before MS analyses based on hydrophobicity using nano-LC on a RP 18 column using a flow rate of 200nL/min. The outlet capillary of the nano-HPLC was coupled with a probot fraction collector (LC-Packings). The LC system delivers the peptides to the mass spectrometer over time. Mass spectrometer first measures the mas-to-charge ratio (m/z) of intact peptides. Selected peptides are fragmented in the collision cell, producing fragment ions in MS1. The unique mass reporter ions (from the TMT tags) are released during fragmentation and are detected in the MS2 spectrum. The intensity of these reporter ions is used to quantify the relative abundance of the peptide across all samples.

### MS data processing

Protein and peptide identification was done with Integrated Proteomics Pipeline (IP2, Bruker Scientific LLC). Tandem mass spectra were extracted from raw files using RawConverter^73^, and searched with ProLuCID^74^ against a database comprising UniProt reviewed (Swiss-Prot) proteome for *Mus musculus* (UP000000589) with Smo (UniProt, P56726) replaced by mSmo-V5-TurboID, Streptavidin (UniProt, P22629) added, and a list of general protein contaminants. The search space included semitryptic peptide specificity with unlimited missed cleavages. Carbamidomethylation (+57.02146 C) and TMT (+229.1629 K and N-terminus) were considered static modifications. Data was searched with 50 ppm precursor ion tolerance and 500 ppm fragment ion tolerance. Identified proteins were filtered using DTASelect2^75^ and utilizing a target-decoy database search strategy^76^ to limit the false discovery rate to 1%, at the spectrum level. A minimum of one peptide per protein and one tryptic end per peptide were required, and precursor delta mass cutoff was fixed at 10 ppm. Statistical model for tryptic peptides (trypstat) was applied. Census2 isobaric-labeling analysis was performed based on the TMT reporter ion intensity using default parameters^77^.

Subsequent data analysis was done in R studio (Supplementary data 4). Sample loading and trimmed mean of M values (TMM) normalization were performed across replicates to have comparable total signal intensities across different replicates. Briefly, channel intensities from three replicates were normalized by calculating the global scaling target, the mean total intensity across all channels from three experiments. For each experiment, the normalization factor is calculated by dividing the global target by the sum of intensities for each channel in the experiment. Each channel’s intensities are then scaled by its respective normalization factor. Next, each channel was subject to TMM normalization to reduce differences caused by variations in sample composition, like highly abundant proteins skewing the total signal. The final normalization dataset, organized by experimental condition and replicate, was subsequently used for downstream statistical analysis.

Differential expression analysis was conducted on the normalized data using an Empirical Bayes moderation approach to stabilize variance estimates. First, comparison conditions were: no Shh/with biotin compared to no Shh/no Biotin and Shh/with biotin to Shh/no biotin. Following this, empirical Bayes moderation was applied to the comparison results using eBayes package in R studio where log-fold changes and adjusted p-values were computed for each protein within the comparisons. All data processing methods and equations can be found in the Supplementary data 5.

Gene enrichment (GO) analysis of Molecular Function was performed using ShinyGO 0.81 (http://bioinformatics.sdstate.edu/go/) for data in Figure 2S.

### Immunofluorescence microscopy of cultured fibroblast

NIH3T3 cells and Smo-V5-TurboID stable cells were grown on poly-D-lysine-coated (Sigma, A003E) coverslips. Once reaching 80–90% confluency, cells were treated with low-serum medium or low-serum medium + 100 nM SAG/5 μM Cyclopamine for the indicated times. Live cells were subsequently incubated with 500 μM Biotin for 15 min at 37 °C, 5% CO2 and washed once with Dulbecco’s PBS before fixation in 4% PFA (Electron Microscopy Sciences, 15713-S). Cells were blocked with blocking buffer (0.2% Triton X-100, 2% Donkey serum in PBS) for 1 h at room temperature. After blocking, cells were incubated with primary antibody at 4°C overnight. Subsequently, cells were incubated with secondary antibodies for 1 h and DAPI for 10 min at room temperature. Cells were mounted in Fluoromount-G (SouthernBiotech, 0100-01). Imaging was performed with Zeiss LSM 880 confocal Laser Scanning Microscope with 100x oil immersion lens or a LEICA DMi8 system with ×63 oil-immersion lens or Leica Mica. Images were processed using FIJI.

### Western Blotting and immunoprecipitation

Standard techniques were used for SDS-PAGE and Western blotting. Cells were first washed with PBS, and then were scraped off from the culture surface in RIPA buffer (1% NP-40, 0.1% SDS, 0.5% sodium deoxycholate, 150 mM NaCl, 25 mM Tris/HCl, pH 7.5, and protease inhibitors). Samples were incubated on ice for 30min. Lysates were cleared by centrifugation (10,000 × *g* at 4 °C for 15 min), and 25 µg protein was separated on 8% SDS-PAGE gels and transferred onto PVDF membranes. After blocking in 5% BSA, membranes were washed and incubated with primary antibodies and secondary antibodies. Finally, proteins were detected with chemiluminescence substrates (Thermo Fisher Scientific, 34076). Quantitation of bands was performed using FIJI.

For Co-immunoprecipitation, HEK293 cells were transfected with YFP-Git1 and Grk2-HA via Calcium phosphate transfection kit (Takara, 631312) according to the manufacturer’s instructions. After 24 h expression, cells were harvested in lysis buffer (20 mM Tris, pH7.5, 150 mM NaCl, 1mM EDTA, 1mM EGTA, 1% Triton X-100, protease inhibitor). Samples were incubated on ice for 30min and centrifuged at 10,000 x g at 4 °C for 15 min. Supernatant was collected and subject to BCA assay for protein concentration. Samples were normalized to the concentration and volume before loading to HA magnetic beads (Thermo Fisher Scientific, 88836) for 30min incubation at room temperature with rotation. Unbound material was removed, and samples were proceeded with western blot (WB) analysis.

### Lentivirus production and concentration

Lentivirus was produced by transfecting plasmids in FUGW lentivirus backbone into 293 T cells using Calcium phosphate transfection kit (Takara Bio, 631312). After 48-h incubation at 37 °C, 5% CO_2_, supernatant was collected and centrifuged at 800 x g for 10min at room temperature. Then, 4 x lentivirus concentrator (40% W/V PEG-8000, 1.2 M NaCl, PBS, pH 7.2) was added to the supernatant and incubator at 4 °C overnight with shaking. Virus was pelleted by centrifugation at 1,600 x g for 60 min at 4 °C and diluted in PBS.

### Quantification and statistical analysis

Cilium length was measured in FIJI. A line was drawn along the fluorescent signal corresponding to the ciliary marker, and the length of this line was defined as the length of the primary cilium. Quantification of ciliary intensity staining was performed in Fiji. The ciliary protein intensity was measured in ImageJ. Briefly, we first outlined the contour of an individual cilium in the channel of cilium staining. After that, the ciliary intensity within this contour was taken for each individual channel of the corresponding protein (L1). Then, the individual cilium contour was manually dragged to the region right next to the cilium, and the intensity within the contour was taken as the background (L2). The final fluorescence intensity for that channel was defined as L = L1/L2. The intensities are reported in Arbitrary units (AU).

Quantification of Gli was performed in FIJI. Briefly, Mean Gray Value was measured for each band using a defined region of interest and an adjacent background value was subtracted. This resulting value for Gli3R was normalized to Gli3FL value in its own lane. This resulting value for Gli1 was normalized to the loading control (GAPDH).

Statistical analysis and graph plotting were performed with GraphPad Prism 8. For analysis of two samples, significance was determined via two-tailed unpaired t test. For more than two samples, significance was determined via one-way ANOVA followed by Tukey’s multiple comparison test for one variable or two-way ANOVA followed by Tukey’s multiple comparison test for two variables. A p value less than 0.05 was considered statistically significant and it denoted as follows: *p < 0.01, ***p < 0.001, ****p < 0.0001, ns, not significant.

